# Transitions in symbiosis: evidence for environmental acquisition & social transmission within a clade of heritable symbionts

**DOI:** 10.1101/2020.07.27.223933

**Authors:** Georgia C Drew, Giles E Budge, Crystal L Frost, Peter Neumann, Stefanos Siozios, Orlando Yañez, Gregory DD Hurst

**Affiliations:** Department of Zoology, University of Oxford, Oxford, UK; School of Natural and Environmental Sciences, Newcastle University, Newcastle upon Tyne, UK; Institute of Integrative Biology, University of Liverpool, Liverpool, UK; Institute of Bee Health, Vetsuisse Faculty, University of Bern, Bern, Switzerland

## Abstract

A dynamic continuum exists from free-living environmental microbes to strict host associated symbionts that are vertically inherited. However, knowledge of the forces that drive transitions in the modes by which symbioses form is lacking. *Arsenophonus* is a diverse clade of bacterial symbionts, comprising reproductive parasites to coevolving obligate mutualists, in which the predominant mode of transmission is vertical. We describe a symbiosis between a member of the genus *Arsenophonus* and the Western honey bee. We then present multiple lines of evidence that this symbiont deviates from a heritable model of transmission. Field sampling uncovered marked spatial and seasonal dynamics in symbiont prevalence, and rapid infection loss events were observed in field colonies and individuals in the laboratory. Fluorescent in-situ hybridization showed *Arsenophonus* localised in the gut, and detection of the bacterium was rare in screens of early honey bee life stages. We directly show horizontal transmission of *Arsenophonus* between bees under varying social conditions. We conclude that honey bees acquire *Arsenophonus* through a combination of environmental exposure and social contacts. Together these findings uncover a key link in the *Arsenophonus* clades trajectory from free-living ancestral life to obligate mutualism, and provide a foundation for studying transitions in symbiotic lifestyle.

## Introduction

Microbes that associate with hosts span a continuum from mutualism to parasitism and employ disparate transmission strategies [1]. Heritable bacterial symbionts, such as *Wolbachia* and *Arsenophonus*, transmit vertically (VT) from parent to offspring [2]. Other microbial symbioses form via horizontal transmission (HT) during host development, by acquisition from environmental reservoirs or via infected conspecifics or other host taxa [3, 4]. In some cases, a combination of these routes may operate, creating a complex transmission landscape [5, 6]. Across this transmission axis lies the impact of the symbiosis on each partner – whether the host and microbe benefit from the interaction, and whether one party requires the other to complete their life cycle. Symbioses vary from being facultative (where one partner does not require the other) to obligate (where there is dependence). Obligacy may be a component of symbiont or host biology, or in some cases both parties are mutually dependent [2, 7, 8].

Clades of heritable symbionts commonly include strains that are both facultative and obligate from the host perspective. In contrast, for the microbe, life is commonly obligately symbiotic, generally lacking replicative or dormant phases outside of host organisms [7]. Despite this dependence, many of these symbionts can also transmit horizontally (inter and intra-specifically) over evolutionary timescales [9–11]. However it is generally accepted that VT predominantly drives population dynamics [12], but see [13]. In contrast, exclusively horizontally acquired symbionts must be transmitted to new hosts via infected con-/hetero-specifics or environmental reservoirs [14, 15]. As a result, symbiotic life for these microbes is commonly facultative [3]. The absence of VT can promote higher rates of partner switching [2, 16] and a weaker association between host and symbiont fitness. Consequently, selection for higher virulence and trajectories towards parasitism are commonly assumed to be favoured more readily in HT symbionts [1, 17, 18], but see [19, 20].

Symbioses thus exist on an evolutionary landscape in which transitions between transmission mode and lifestyles occur. However, study of the drivers and impact of transitions in transmission mode is inhibited by a lack of clades in which different lifestyles co-occur among members. Clades encapsulating diverse transmission biology enable understanding of both the ecological and evolutionary mechanisms driving transitions, and the consequences of these for processes such as genome evolution [21–24]. For instance, the emergence of a virulent horizontally transmitted vertebrate pathogen (*Coxiella burnetti*) from a clade of maternally inherited tick endosymbionts [22] allows insight into how infectious transmission can emerge from a heritable, obligate clade. Likewise, the presence of an opportunistic human infective *Sodalis* provides an important comparator for understanding the evolution of symbiosis, through comparison to insect-associated mutualistic lineages of *Sodalis* [21].

As a monophyletic clade of heritable Enterobacteriaceae, *Arsenophonus* provides a valuable base for exploring the evolution of a heritable lifestyle [25], due to its wide host distribution (est. 5% of all arthropod species) [26] and diversity in symbiotic lifestyle [27]. Members of the clade include reproductive parasites [28], facultative mutualists [29, 30] and highly coevolved obligate endosymbionts undergoing genomic decay [31–33]. Despite this diversity, all strains characterised to date are vertically transmitted. While frequent horizontal transmission over ecological timescales has been shown for a number of strains, including the male-killer *Arsenophonus nasoniae* [13, 34, 35] and two species of insect-vectored phytopathogens (*Arsenophonus phytopathogenicus* and *Phlomobacter fragariae*) [36–38], this route occurs alongside vertical transmission. As a result, *Arsenophonus* is commonly reported from arthropod screening efforts and presumed heritable without further characterisation.

An economically important eusocial host, the Western honey bee (*Apis mellifera*), has previously been associated with *Arsenophonus* [39–43] and infection has been linked to poor health outcomes [44] including colony collapse disorder [45]. Whilst this interaction has attracted interest from the community, basic information on the epidemiology and transmission of *Arsenophonus* in honey bee populations is nevertheless lacking. Symbionts in eusocial hosts are exposed to markedly different selection pressures from those in solitary species. This difference arises from a higher density of hosts, greater host relatedness and a homeostatic nest environment as well as reproductive division of labour, overlapping generations and cooperative brood care [46, 47]. Specialised social behaviours [48] additionally foster the transmission of microbes by direct contact, such as via proctodeal (anus – mouth feeding) and stomodeal trophallaxis (mouth-mouth feeding) [49]. Thus, host sociality is an important driver of symbiont phenotypes [47, 49, 50], and indeed interesting heterogeneities in symbiont infections are emerging based on caste, sex [51–55] and degree of sociality [50]. Despite this, the potential effects of host sociality on symbiont ecology and evolution, including important phenotypes such as transmission, remain largely unexplored.

This study focuses on characterising the ecology and transmission of *Arsenophonus* from a eusocial host. We use a phylogenomic approach to show robust placement of the strain within the *Arsenophonus* clade. We then present data tracking the prevalence of the symbiont in honey bee colonies over space and time, to search for indicators of stable maintenance (implying VT) or changes in prevalence (implying infectious transmission epidemics). These data are complemented by FISH analysis highlighting tissue interactions with honey bees. Finally, the capacity for vertical transmission is assessed directly by screening for *Arsenophonus* across host life history, and the ability to horizontally transmit intra-specifically is investigated under differing social conditions. These data demonstrate the presence of an infectiously transmitting *Arsenophonus* without vertical transmission, highlighting transitions in life history within this important genus of insect-associated microbes.

## Methods

### Phylogenomic position of *Arsenophonus* from honey bees

To assess the relatedness of the *Arsenophonus* strain associated with honey bees to other *Arsenophonus*, a phylogenomic approach was adopted. To this end, a draft genome was assembled (bioproject accession: PRJEB39047) using an Illumina paired end shotgun library using reads obtained in a previous study (see Gauthier et al., 2015). In total, 53 core ribosomal protein coding genes were extracted and compared to available *Arsenophonus* genomes. Relatedness of strains was estimated by Bayesian inference on the bases of the concatenated alignment of the 53 ribosomal protein dataset, completed using Phylobayes-MPI [57] and the CAT-GTR model. Two independent chains were run in parallel for over 25,000 cycles each until convergence occurred (maxdiff < 0.1).

### Spatial and seasonal dynamics of *Arsenophonus* in honey bee colonies

To monitor *Arsenophonus* prevalence over time and space, adult workers were collected from the outer frames of 159 colonies stemming from 45 apiaries in 10 counties across England. Colonies were sampled from April to November during 2014 to 2018, and repeated screening of some colonies resulted in a total of 230 sampling events. Apiaries are defined here as a collection of colonies that are colocalised within a small area, such as a field. Bees were preserved in 70% EtOH at −20°C until DNA extraction. For each colony, posterior legs were pulled from workers (n=12), pooled in groups of four and exposed to UV light for 10 min to cross link DNA from surface microbes. Legs were used as this tissue was found to be a reliable marker of *Arsenophonus* association in bees and did not inhibit downstream PCR assays when DNA was extracted using a high throughput Chelex protocol [58]. Extraction quality was verified by amplification of host DNA (EF1-α) [59] and the presence of *Arsenophonus* spp. established by PCR assays targeting *fbaA* (adapted from Duron et al. 2010, see SI) and Sanger sequencing of the product to determine broad identity of the *Arsenophonus* strains. Sensitivity of PCR assays was established through serial dilution, and was robust over two orders of magnitude (see SI).

To determine if *Arsenophonus* is lost from honey bee colonies during over-wintering, the status of 25 colonies (from 7 apiaries) was tracked from autumn to spring at a greater depth than described above. In the autumn, 15 honey bee workers were screened from each colony (3 individuals pooled per extraction) to determine *Arsenophonus* prevalence. If 80% of extractions (i.e. 4 of 5) were positive for *Arsenophonus* the colony was included in the infected cohort (group A, n=19). The uninfected cohort (group B, n=6) comprised of colonies where all extractions returned negative. Colonies were left to overwinter, and the sampling process was repeated in the spring for colonies that overwintered successfully. To determine infection status in spring a total of 24 bees were screened per colony (8 pools of three bees). DNA was extracted from whole bees (heads removed) by Promega Wizard® purification, with PCR detection of *Arsenophonus* (see SI).

### Localisation of *Arsenophonus* within the gut

To visualise *Arsenophonus* using fluorescence in situ hybridization (FISH), whole guts were dissected from live honey bee workers and placed into Carnoy’s fixative (60% EtOH, 30% chloroform, 10% acetic acid) for 24–48hrs, washed with 100% EtOH (x 3) and incubated with hybridization buffer (20Mm Tris-HCL, 0.9M NaCl, 0.01% SDS, 30% formamide, 100 pmol/ml probe, see SI for detail) at room temperature in the dark (∼15 hrs). The symbiont was targeted using an *Arsenophonus* specific probe (TCATGACCACAACCTCCA) [60] with a 5’ Alexa Fluor® 647 fluorochrome. Tissues were washed with pre-heated (∼48°C) buffer (20Mm Tris-HCL, 0.9M NaCl, 0.01% SDS, 0.05M EDTA) x 3 for 10 min (room temperature, dark) and mounted on glass slides with DAPI containing ProLong Diamond anti-fade (Fisher Scientific). Tissues cured for 24 hrs before visualisation by confocal microscopy (ZEISS LSM 880 with x 40 objective lens), multiple optical sections were assembled into Z-stacks under maximum intensity settings using ImageJ [61]. Gut tissue of honey bees from colonies not associated with *Arsenophonus* underwent the same process to function as negative controls. In addition to gut imaging, faecal samples were collected from workers (n=22) isolated in sterile petri dishes. Faecal material was removed by aspiration with molecular grade H20 (∼50 μl) and extracted by Promega Wizard® purification (see SI). Material was then tested for *Arsenophonus* DNA through PCR assay.

### Assessing the heritability of *Arsenophonus* and acquisition from infected conspecifics

To determine heritability of *Arsenophonus* in honey bees, infection was assessed across life history stages. Frames of honey bee worker brood were removed from managed field colonies (N= 8, A+ = 6, A-=2) where *Arsenophonus* status had previously been determined with high confidence. Eggs (n= 9), larvae (n=67), pupae (n=49), newly emerged workers (NEWs) (n=36), adult workers (n=45), and where possible drones (n=22), were collected concurrently. DNA was extracted individually from eggs and early stage larvae using a Qiagen ® DNeasy blood & tissue kit, all other life stages were extracted by Promega Wizard® purification. Molecular detection was completed as previously.

To assess if infections were maintained in the absence of the colony and foraging environment, infected individuals were detained in the laboratory and their *Arsenophonus* status tracked over 15 days. Adult worker bees were collected from two colonies (total N=76, Colony A=36, Colony B=40) with a high prevalence of *Arsenophonus* in 15 adult bees (> 90%) or one control colony where none of 15 adult bees tested positive (N=32) (based on *Arsenophonus* detection of individual bees, see SI). On entering the laboratory (day 1) worker bees were cooled to 4°C and the *Arsenophonus* status of all individuals was established by removal of posterior tarsus tissue (‘leg snip’) and PCR assays for symbiont presence (see SI). Additional bees (‘no leg snip control’, N=40) from the infected source colonies were not subjected to leg snips to establish if this method biased results. Individuals were marked with queen paint and maintained in groups of four. To track the maintenance of *Arsenophonus* over time, individuals were randomly selected from each group and culled at days 4, 8 and 15. Individual *Arsenophonus* status was determined using the same methodology as day 1, but using opposing tarsus tissue.

To assess the capacity for horizontal acquisition of the symbiont, adult workers (donors) were taken from colonies (N = 6) with a high prevalence of *Arsenophonus* in 15 adult bees (> 85% infected, see SI) and mixed in 340ml pots with uninfected newly emerged workers (recipients). Each pot contained 10 donors and 5 recipients. Two transmission treatments were established, each with 10 replicates. In the ‘general contact’ treatment, bees were allowed to freely contact one another within the pot and food (50% sucrose solution) was openly available. In the ‘trophallaxis’ treatment, recipients with no direct access to food were separated from donors (with food) by fine mesh, forcing infected bees to feed uninfected individuals by trophallaxis, but preventing other social contact. Contacts were allowed for 5 days, after which the fate (dead or alive) of all individuals was recorded, i.e. those that had died during the course of the experiment (prior to final cull) were labelled dead and analysed separately. In each case, experiments were compared to controls in which uninfected bees were mixed with recipient bees. DNA was extracted individually from whole bees (test; n=300, control; n=165) by Promega Wizard® purification.

### Statistical methods

For statistical analyses, generalized linear models (GLM) and mixed models (GLMM) were fitted in R version 3.3.1 [62] with binomial error distributions and logit link function, using the packages *glm* and *lme4* [63]. Minimum adequate models (MAM) were selected by Akaike Information Criteria (AIC) [64, 65] and likelihood ratio tests (LRT), with the latter being used to assess the significance of fixed and random effects [66]. Overdispersion was assessed using the Blemco package [62]. See supplementary information for tables detailing selection of statistical models.

## Results

### Phylogenomic position of the honey bee *Arsenophonus* in the wider clade

Phylogenomic analysis based on 53 ribosomal protein sequences unambiguously placed the honey bee associated strain within the *Arsenophonus* clade (Figure 1) alongside known heritable symbionts. The strain was established as a sister strain, with strong support, to *Arsenophonus* strains infecting other hymenopteran hosts (parasitoid wasps), including the male-killing reproductive parasite *A*. *nasoniae*.

**Fig. 1.**
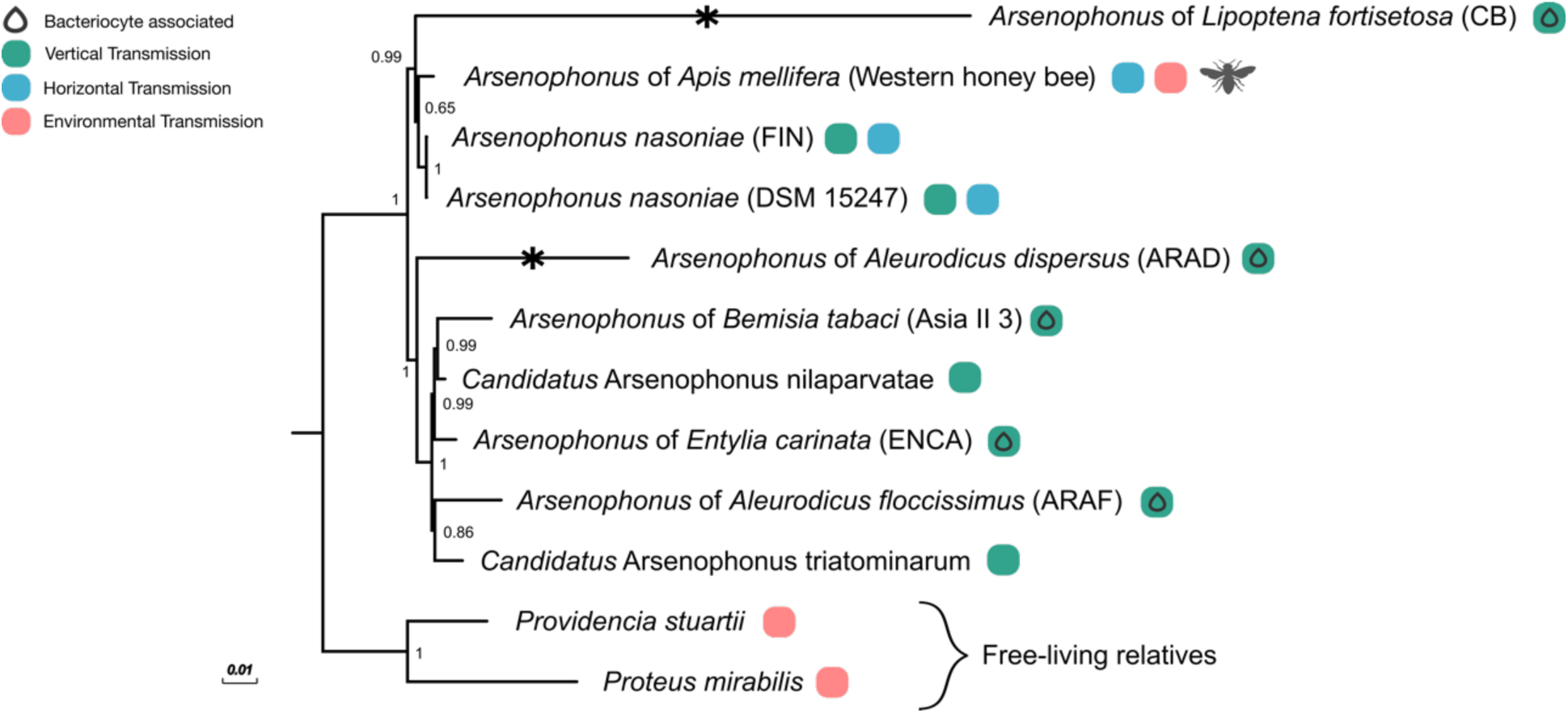
Transmission mode and lifestyle diversity in the *Arsenophonus* clade. Phylogenomic position of the *Arsenophonus* associated with honey bees (this study) among *Arsenophonus* spp. with available genomes. Bacterial species names are given, if formally recognized, else the associated insect host is noted. Symbols indicate known bacteriocyte associations and transmission modes, vertical transmission (green), horizontal transmission (blue), if an environmental transmission route is additionally inferred this is indicated in pink. Analysis based on 53 ribosomal proteins, support for Bayesian inference (posterior probabilities) is shown at nodes.

### Spatial and seasonal dynamics of *Arsenophonus* in honey bee colonies

*Arsenophonus* was found associated with 38.9% (95% CI: 32.5-45.6%, N=159 colonies) of honey bee colonies across the UK. Infections were spatially widespread and identifiable in all 10 county regions tested (Figure 2.A). County region was not a significant predictor of *Arsenophonus* infection (GLMM, LRT: X^2^=0, *df=*1, *P=*1), but spatial patterns were evident at a local scale, with apiary emerging as an important covariate, with colonies within an apiary having correlated infection status (GLMM, LRT: X^2^=13.8, *df=* 1, *P=*0.002**). All *Arsenophonus* detected had identical sequence at the *fbaA* locus (N=159).

**Fig. 2.**
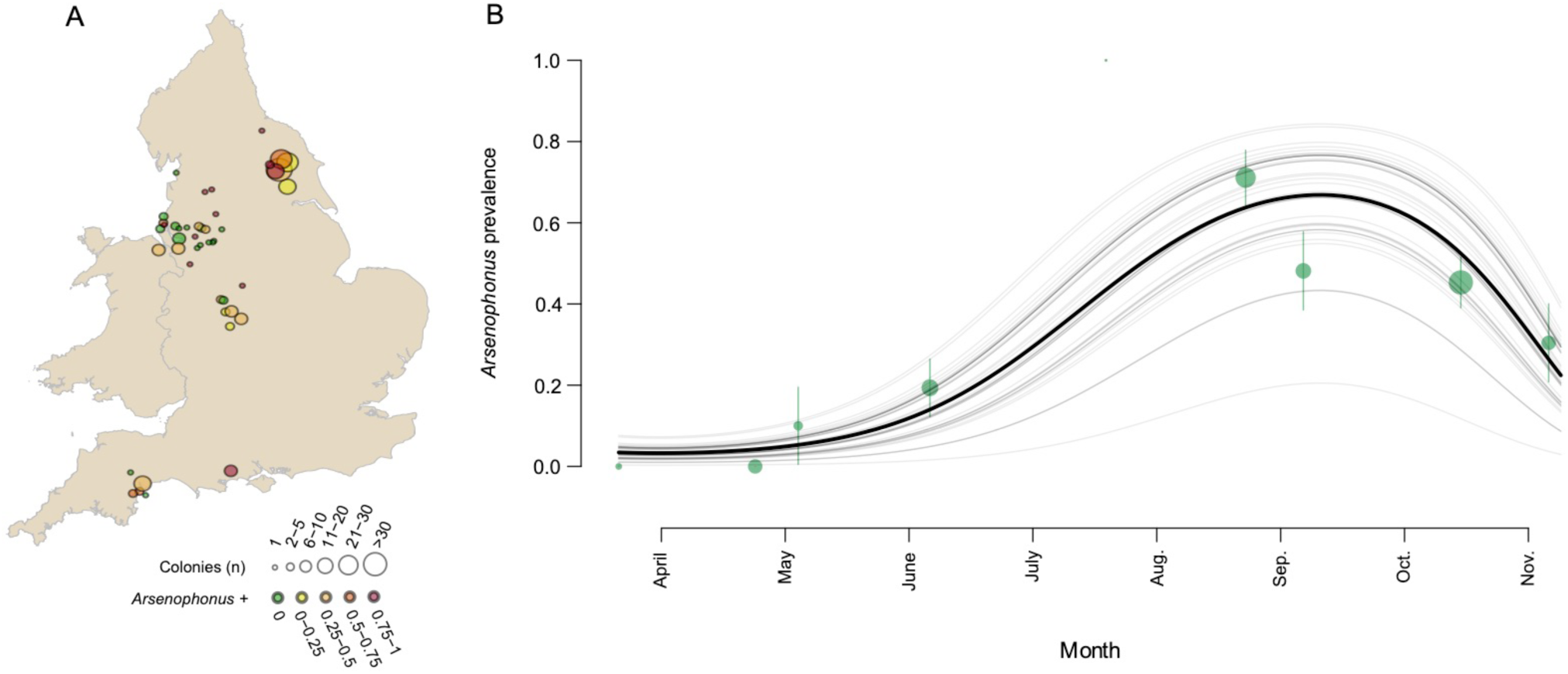
Spatial and seasonal dynamics in the *Arsenophonus –* honey bee association. (A) Circles show the approximate locations of 45 honey bee apiaries (colonies colocalised within a small area) sampled for *Arsenophonus* across England (2014 – 2018). Circle size reflects the number of colonies sampled within an apiary location and colour indicates the proportion of colonies associated with *Arsenophonus*. (B) *Arsenophonus* prevalence in honey bee colonies from spring to autumn modelled using 229 binomial observations of *Arsenophonus* status from 159 colonies (2014 – 2018). An overall prediction for all apiaries (black) and individual predictions for each apiary (grey, n=45) are shown. Green circles represent the mean prevalence of *Arsenophonus* by month at the mean monthly sampling date, size is proportional to the square root of the number of colony samplings. Error bars represent binomial CI.

Strong seasonal dynamics in *Arsenophonus* infections (Figure 2.B) were evident, with prevalence of the bacterium changing in a non-linear fashion during the main foraging period of the host (March to October). *Arsenophonus* prevalence is lowest during the spring (2.63% of colonies, 95% CI: 0.0670-13.8%, N=38) with the first infected colony detected in May. Prevalence continues to rise, reaching a peak in late summer (August: 71% of colonies, 95% CI: 55.7-83.6, N=45), before dropping into Autumn (43.5% of colonies, 95% CI: 34.3-53.0, N=115). Significant temporal variation by year was observed for *Arsenophonus* prevalence (GLMM, LRT: X^2^=3.87, *df=*1, *P=*0.049*). See SI Table 1 for a full summary.

In temperate environments, overwintering conditions represent a distinct state for honey bees, with foraging ceasing temporarily and survival dependent on stored colony resources. The *Arsenophonus* infection status of 25 honey bee colonies was tracked from autumn to spring (Figure 3). Of the 23 colonies that survived winter, all of those infected with *Arsenophonus* in the autumn (Group A, n=17) had lost the bacterium by spring. Of the colonies where *Arsenophonus* was not detected in the autumn (Group B, n=6), five remained uninfected in spring, and one case gained *Arsenophonus*. Confidence in this newly positive colony is high, as all pools tested (N=8 pools of 3 bees) were positive for *Arsenophonus*. Notably, this colony was sampled latest in the spring season.

**Fig. 3.**
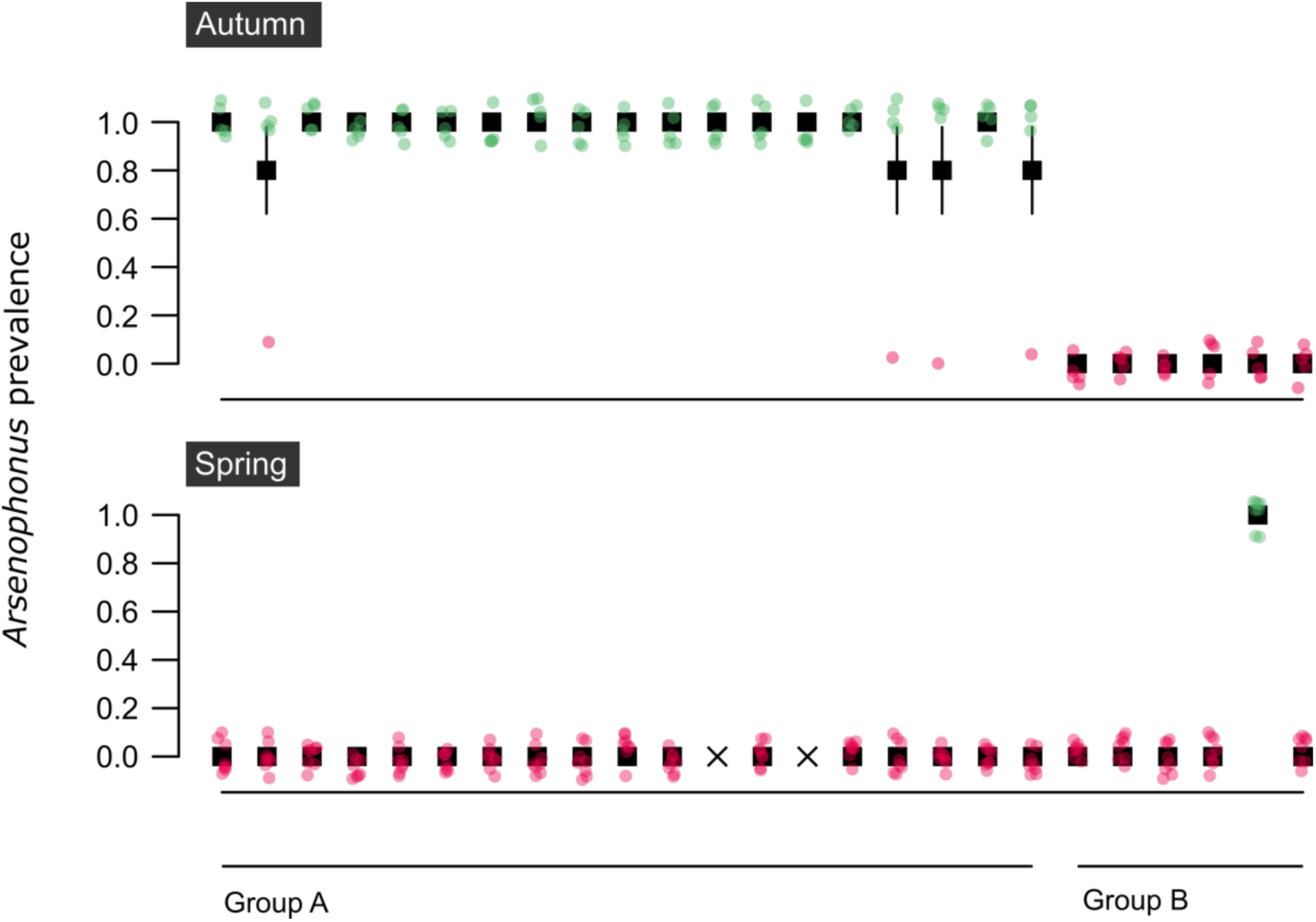
Overwintering loss of *Arsenophonus* at a colony level. Group A colonies (n = 19) were infected with *Arsenophonus* in autumn but the bacterium was undetectable by the following spring. Group B colonies (n = 6) were uninfected with *Arsenophonus* in autumn, one colony gained *Arsenophonus* association by spring. Colonies that did not survive the winter (n=2) are denoted by an X. Mean *Arsenophonus* prevalence by colony is shown. Note, fewer bees were sampled in Autumn (n=15 bees per colony, pooled in groups of 3) than spring (n=25 bees per colony, pooled in groups of 3). Pink dots (0 = *Arsenophonus -*) and green dots (1 = *Arsenophonus* +) show the raw binomial data jittered. Error bars indicate binomial SE.

### Localisation of *Arsenophonus* within the gut

Previous work using metagenomic analysis has reported *Arsenophonus* in the gut [40, 41], on the cuticular surface [39] and in haemolymph [56]. To obtain confirmation of symbiosis, targeted imaging of gut tissue from infected honey bees was conducted. This analysis showed aggregations of *Arsenophonus* (coloured red) in the midgut (Figure 4) sufficiently large to imply colonisation and replication. Control images of uninfected honey bee tissue suggested neither autofluorescence nor inadequate probe removal contributed to artefactual *Arsenophonus* visualisation (see SI Figure 1). *Arsenophonus* was also detected by PCR assay in 59.1% of faeces from infected bees (95% CI: 36.4-79.3) suggesting the bacterium may be shed from the gut, however viability of the symbiont was not confirmed.

**Fig. 4.**
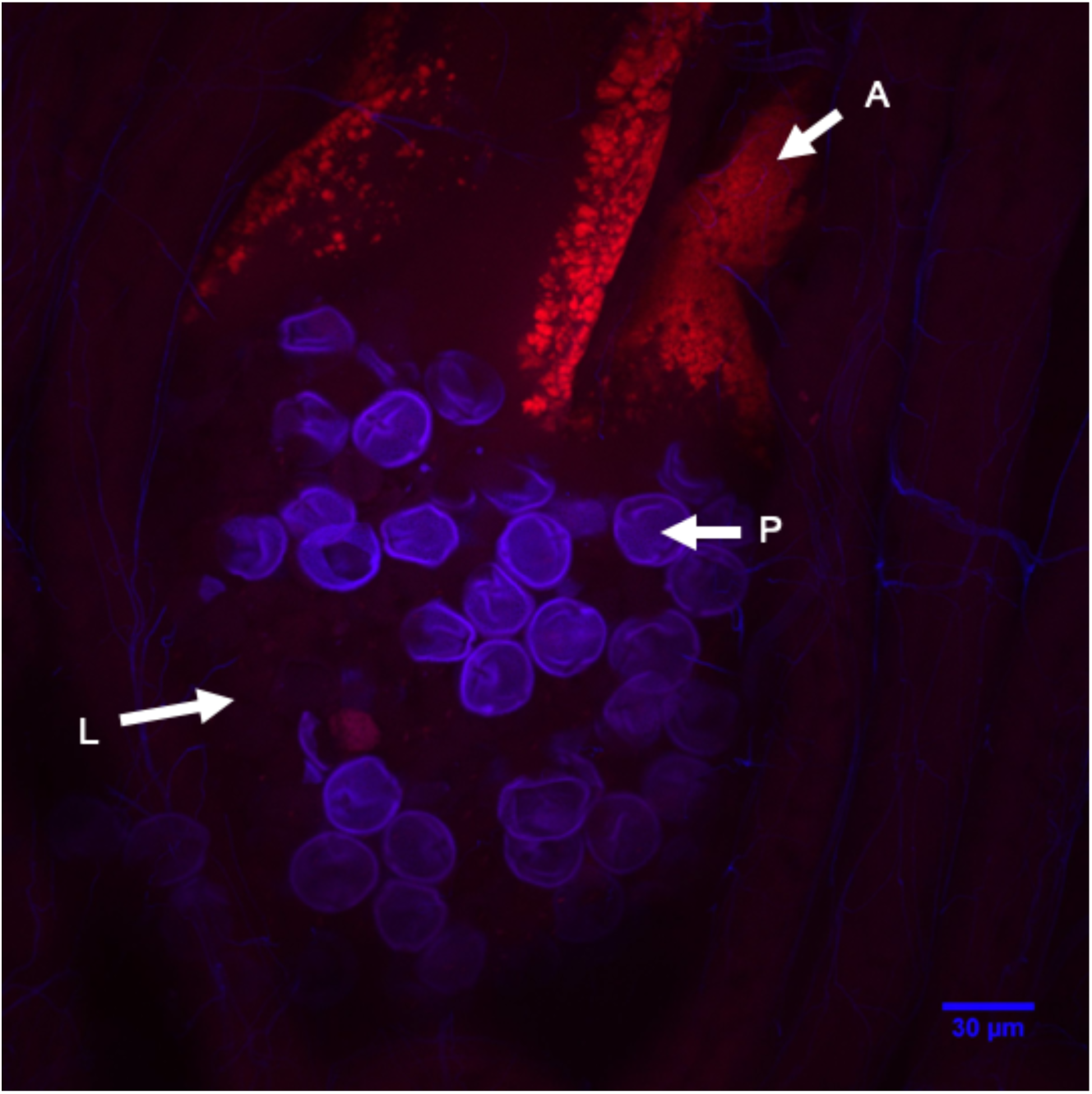
Localisation of *Arsenophonus* bacteria in the midgut of a worker honey bee. Confocal microscopic image of a whole honey bee gut mount with an *Arsenophonus* specific Alexa Fluor® 647 labelled probe (red fluorescence) and DAPI counter-staining (blue fluorescence). Within the midgut lumen (L) pollen grains (P) and hybridization of the *Arsenophonus* specific probe (A) are visible. The image is a composite Z-stack comprised of 32 optical slices (ZEISS LSM 880) assembled in ImageJ under maximum intensity. For confocal images of whole gut mounts from honey bees uninfected with *Arsenophonus* see SI Figure 1.

### Assessing the heritability of *Arsenophonus* and acquisition from infected conspecifics

To date, all characterised *Arsenophonus* strains show vertical transmission in their arthropod hosts. To assess if the honey bee strain conforms to this transmission strategy, the presence of *Arsenophonus* was assessed across host life history (Figure 5). There was no evidence of transovarial transmission of *Arsenophonus* with eggs consistently testing negative, corroborating previous findings in honey bees [43]. In other early life stages *Arsenophonus* was detected only at low frequencies, with 5.97% of larvae (95% CI: 1.65-14.6, *P* < 0.001***), 4.081% of pupae (95% CI: 0.498-14.0, *P* < 0.001***) and 2.78% of newly emerged workers (NEWs) (95% CI: 0.07-14.5, *P* < 0.001***) testing positive for *Arsenophonus*. In contrast, adult life stages showed high incidences of *Arsenophonus*, with the bacterium detectable in 54.5% of drones (95% CI: 32.2-75.6, *P=*0.0198^*^) and 82.2% of workers (95% CI: 67.9-91.0).

**Fig. 5.**
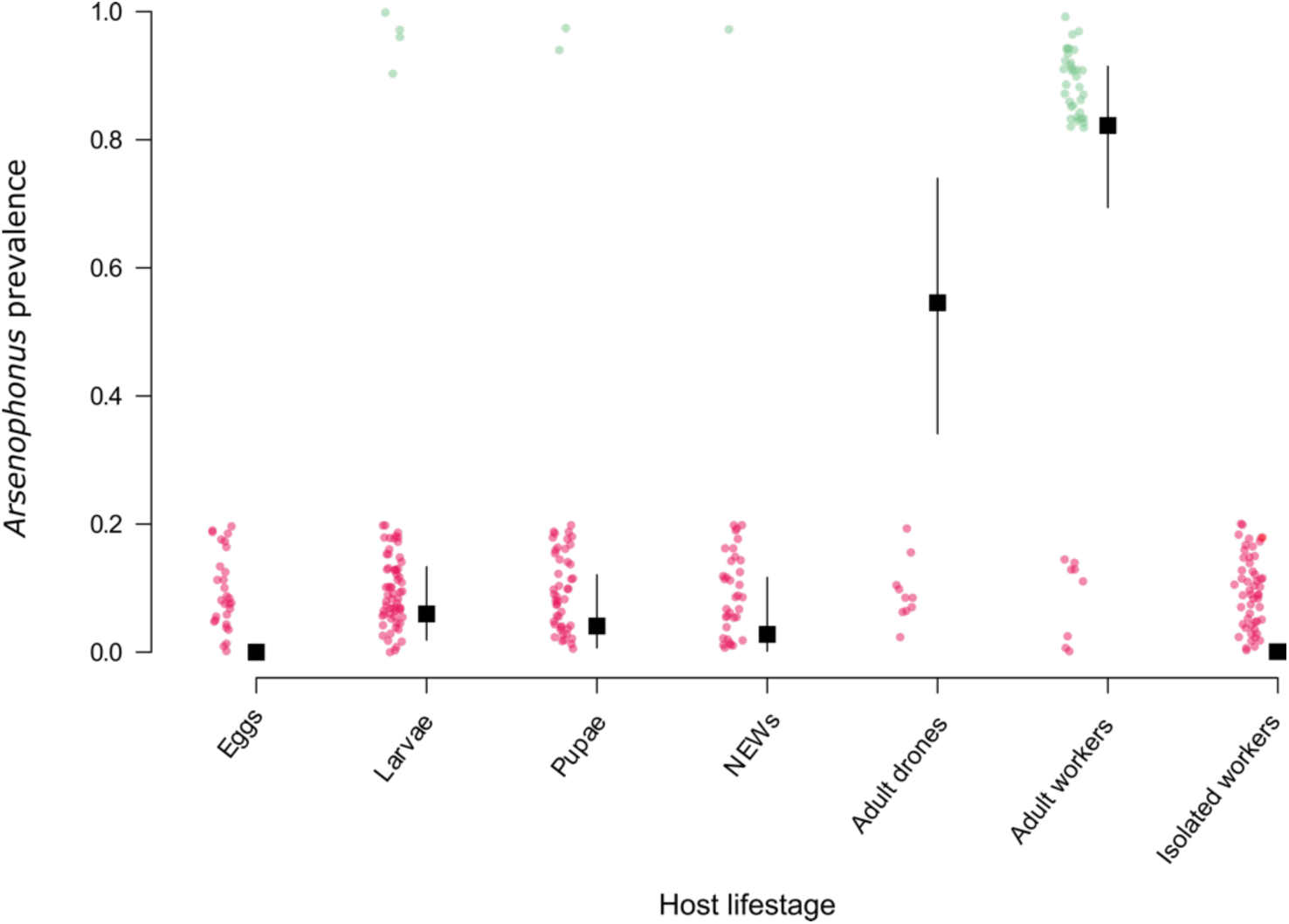
Vertical transmission is largely absent in the *Arsenophonus* – honey bee association. Samples from across the host life history were taken from colonies (n = 6) identified as *Arsenophonus* positive (based on workers) and screened for *Arsenophonus*. Eggs, larvae, pupae and newly emerged workers (NEWs) were all of worker caste. Isolated workers were removed from colonies as NEWs and allowed access only to other NEWs before screening for *Arsenophonus* at forager age (25 days ± 2). Binomial GLM predictions are shown with 95% CI. Pink dots (0 = *Arsenophonus -*) and green dots (1 = *Arsenophonus* +) show the raw binomial data jittered.

The maintenance of *Arsenophonus* in worker bees in the absence of sources of infection was then tested. Honey bees were removed from two infected colonies (colony A and colony B) and maintained in the laboratory for 15 days with access only to sterile food (Figure 6). At day 0 all individuals in the leg snip group were confirmed positive for *Arsenophonus*. Individuals in the ‘no snip control’ group were presumed, from identical colony origin, to be infected at a similar prevalence. There was no significant effect of treatment (leg snip *or* no snip) on the proportion of honey bees infected with *Arsenophonus* at each time point (LRT: X^2^= 0.253, *df=*1, *P=*0.615), suggesting the leg snip did not impact our results. Overall, the proportion of honey bees infected with *Arsenophonus* decreased significantly over time (*P* < 0.001***) indicating loss of the bacterium. However, variation was evident at a colony level, and a significant effect of colony was observed (LRT: X^2^=20.2, *df=*1, *P* < 0.001***). Colony B lost *Arsenophonus* at a significantly faster rate (*P* < 0.001***), with fewer than 20% of individuals infected by day 15. Control honey bees taken from uninfected colonies maintained 0% infection throughout the study. See SI Table 2 for a full summary.

**Fig. 6.**
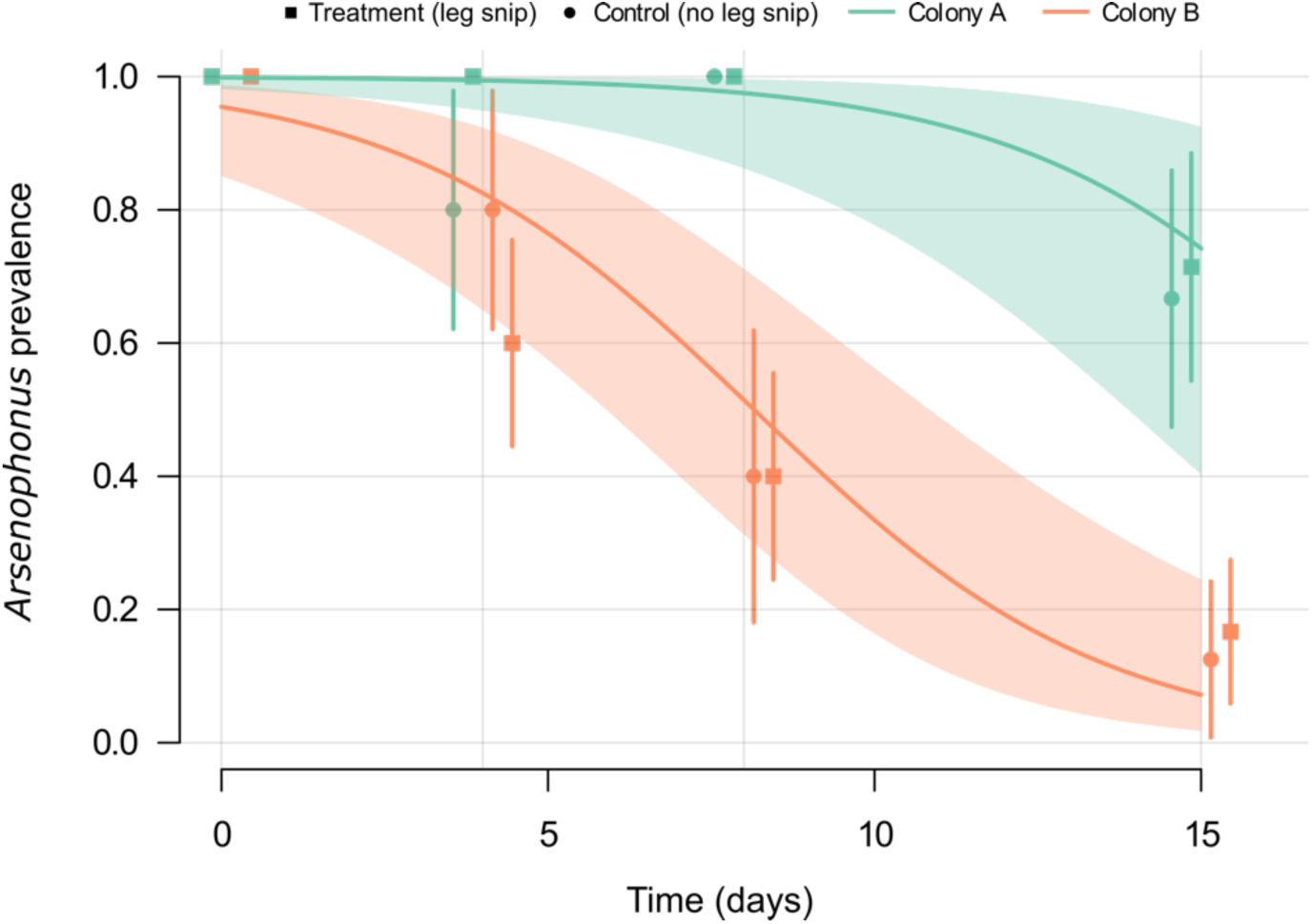
Loss of *Arsenophonus* in individuals removed from the colony and foraging environment. Binomial GLMM predictions (with 95% CI) show the proportion of individual honey bees that remain infected with *Arsenophonus* over time when removed from colonies and detained under lab conditions. The initial *Arsenophonus* status of each bee was determined by a leg snip (treatment group). An additional group of bees from the same colonies did not undergo leg snip (control group), and thus *Arsenophonus* starting prevalence was unknown. Overall predictions are shown for colony A (green) and colony B (orange), and the proportion of honeybees infected at day 0, 4, 8 and 15 is plotted independently for each colony and treatment.

Horizontal acquisition of *Arsenophonus* was tested in a separate experiment, where uninfected recipient bees were exposed to infected donor bees, either through trophallaxis or general contact. Acquisition of *Arsenophonus* among recipient bees was observed under both of these social conditions (Figure 7.A). The rate of *Arsenophonus* gain for recipients was higher under general contact exposure (recipients, 40.0%, 95% CI: 26.4-54.8) compared to trophallaxis (recipients, 22.0%, 95% CI: 11.5-36.0) and social context had a significant effect in the model (GLMM, LRT: X^2^= 5.22, *df =* 1, *P=*0.0223*).

**Fig. 7.**
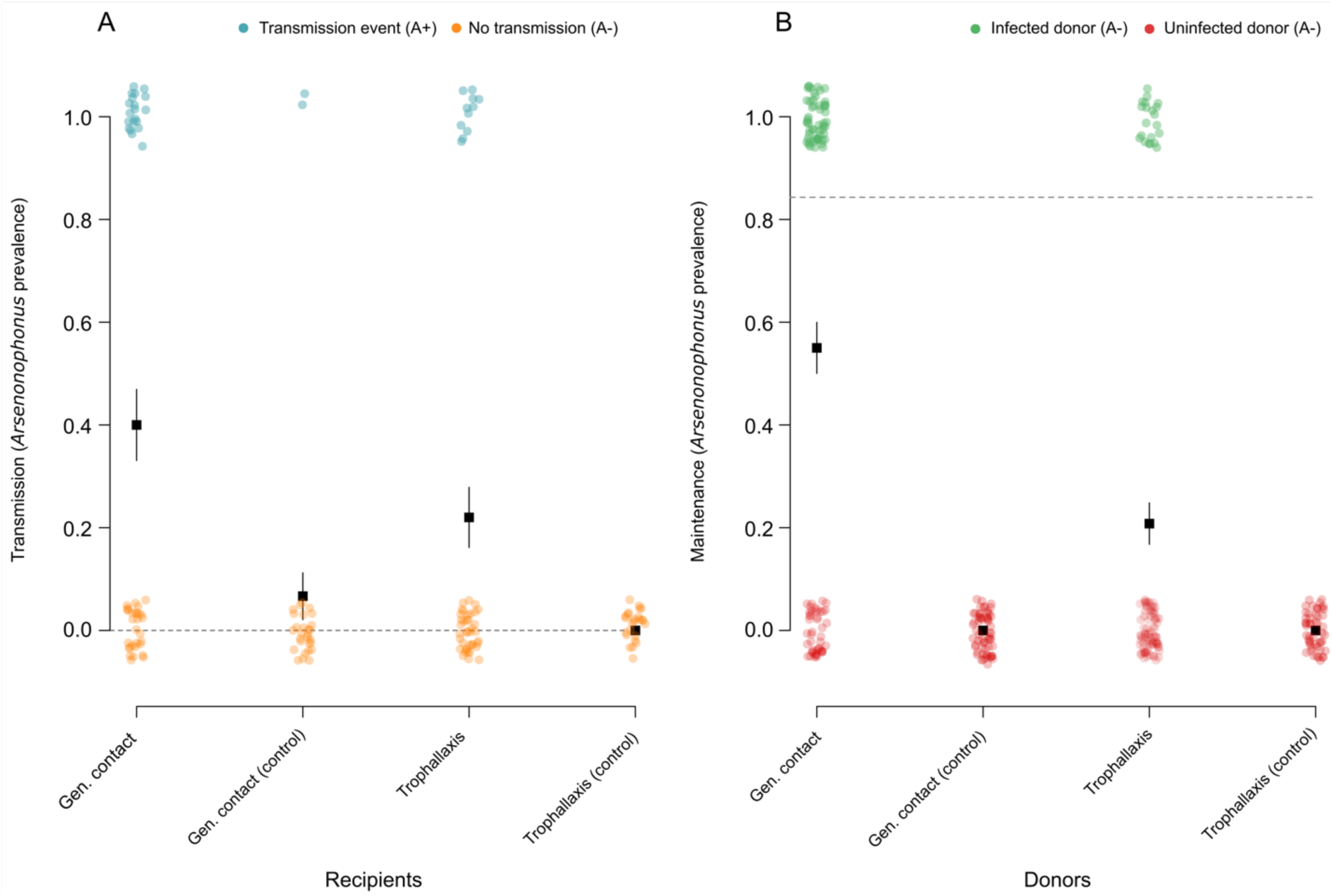
Horizontal transmission and maintenance of *Arsenophonus* in honey bees under two social conditions. (A) Uninfected (recipient) bees were mixed with (B) infected (donor) bees and allowed either general contact or contact via trophallaxis only. For control groups, recipients were mixed with uninfected bees. Dotted lines indicate the prevalence of *Arsenophonus* in recipients and donors at the start of the transmission period. Note, all control bees started uninfected (*Arsenophous* prevalence = 0%). After 5 days of social interaction the *Arsenophonus* status of recipients (A) and donors (B) is shown. Transmission to recipients occurred under general contact and trophallaxis, but at varying levels. Recipients mixed with uninfected (control) bees did not acquire *Arsenophonus*, with the exception of two individuals in the general contact treatment. Coloured dots show raw binomial data jittered (0 = *Arsenophonus -*, 1 = *Arsenophonus* +). Each group was replicated (n = 10), with 5 donor bees and 10 recipient bees per replicate.

Within this experiment, a subset of donors lost the infection from the 85% prevalence starting threshold (Figure 7.B), paralleling the previous observation that infection was not stable under laboratory isolation. Horizontal acquisition between donor and recipient bees contrasted to control pots where the same contact was with uninfected ‘donor’ bees. Here, recipient controls under trophallaxis remained uninfected (control recipients, 0%, 95% CI: 0-13.7), however two control recipients under general contact tested positive for *Arsenophonus* (control recipients, 6.66%, 95% CI: 0.818-22.1). The fate of individuals (live/dead at end of transmission period) emerged as an important predictor of infection status (GLMM, LRT: X^2^=7.64, *df =*1, *P=*0.00571**), with individuals that had died during the study more commonly associated with *Arsenophonus* infection (Figure 8). See SI Table 3 for a full summary.

**Fig. 8.**
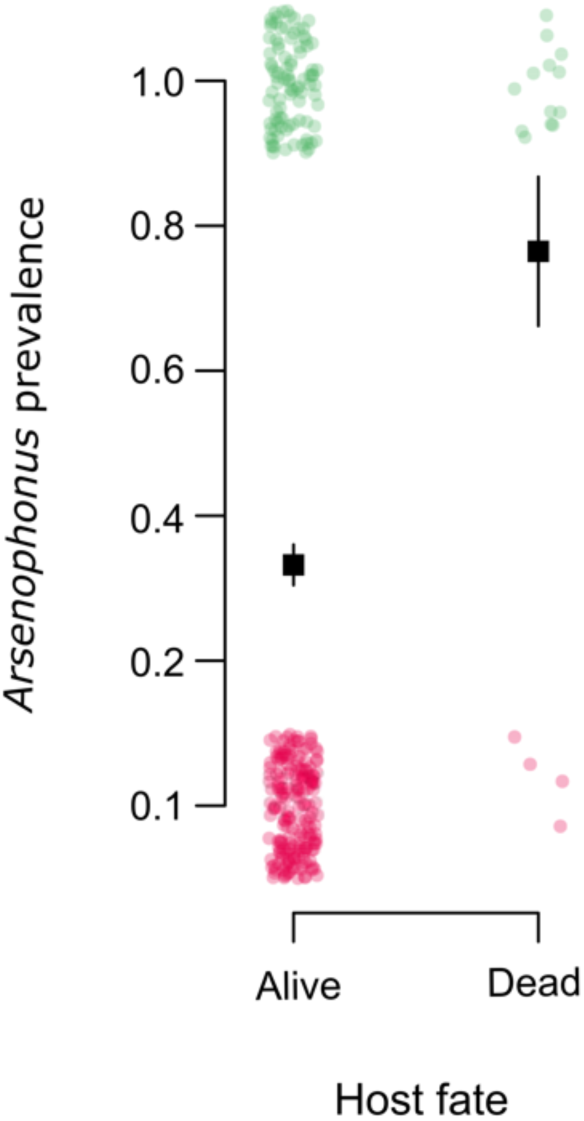
Association between *Arsenophonus* status and honey bee fate. *Arsenophonus* prevalence was higher among individuals that died (n=17), compared to those that remained alive (n=283), during the course of the horizontal transmission study. Pink dots (0 = individual *Arsenophonus -*) and green dots (1 = individual *Arsenophonus* +). Errors bars indicate binomial SE.

## Discussion

The diversity of existing symbiotic interactions represent the result of historical evolutionary transitions between lifestyles. Here, we investigate the symbiotic lifestyle of *Arsenophonus* associated with honey bees. A significant contingent of the *Arsenophonus* clade is in the latter stages of an evolutionary trajectory towards obligate intracellular life [31–33], and until now all characterised members of the genus demonstrate a heritable lifestyle, at least to some degree [26, 27]. We here show that *Arsenophonus* associated with honey bees deviates significantly from this heritable model, instead demonstrating strong seasonal patterns and a dynamic association that appears to be driven by social transmission with an additional, unidentified, environmental reservoir. These results shed new light on evolutionary transitions and symbiotic diversity within a clade of heritable arthropod symbionts.

We have identified the *Arsenophonus* - honey bee strain as the first within the clade to show little evidence of heritability, with evidence for this coming from multiple sources. First, the absence of the bacterium in eggs of infected colonies corroborates previous findings that *Arsenophonus* is not transovarially transmitted in honey bees [43], while very low prevalence in early host life stages provides new evidence that no alternative routes of VT are operating in the association. Secondly, the pronounced seasonal dynamics we observed for *Arsenophonus* are not consistent with the epidemiology of heritable symbionts, whose infection prevalence generally remains relatively static within a host generation though may vary dynamically between generations [67]. Within these dynamics, colonies appear to lose *Arsenophonus* over the winter, or the association declines to a level undetectable by our survey. Finally, experiments show first that honey bee workers with *Arsenophonus* infection lose it under laboratory conditions under social isolation, and also that social contact can result in acquisition of infection. Overall, *Arsenophonus* presents as a horizontally acquired infection of honey bees rather than a persistent vertically transmitted one.

This acquisition-loss cycle occurs for a bacterium that is otherwise nested into a clade of heritable microbes, with phylogenomic analysis indicating *Arsenophonus* from bees lies in the same subclade as the male-killing microbe, *Arsenophonus nasoniae*. Whilst predominantly associated with the parasitic *Nasonia* wasp, *A*. *nasoniae* is an extracellular symbiont that also has phases outside of the wasp host [68], is culturable [28], and is also maintained by a combination of vertical and infectious transmission [13]. *Arsenophonus nasoniae* and the bee *Arsenophonus* also share the capacity to establish through the gut [68], indicating this pathway is preserved in the clade. The genome degradation process that occurs in the obligate symbionts in the genus imply that the bee *Arsenophonus/A*.*nasonaie* lineage within the clade has retained the extracellular and environmental growth capacities found in a free-living ancestor. The alternate model - reacquisition of this capacity – is less probable as this would require the re-establishment of multiple genetic systems allowing growth outside of a host environment. This scenario is not, however, impossible, as previous work on *Coxiella* indicated the striking emergence of an infectious vertebrate pathogen from a clade of heritable, and sometimes obligate, endosymbionts of arthropods [22].

Our data thus indicate that the clade *Arsenophonus* contains more transmission diversity than previously considered. Knowledge of the selective forces that drive the emergence of symbiotic diversity is imperative for understanding important transitions such as the emergence of pathogenic agents from commensal partners, and vice versa [21, 22, 69].

Enterobacteriaceae genera appear to be well adapted for transitioning between ecological and symbiotic niches (Sachs et al., 2011; Walterson and Stavrinides, 2015); for example, *Pantoea, Sodalis* and *Serratia* include notable symbionts of insects [72–74] but are largely comprised of representatives from soil, plants and clinical settings [21, 71, 74]. Likewise, the genus *Photorhabdus* includes both defensive symbionts and environmentally acquired pathogens of invertebrates [75, 76], but also a primary pathogen of humans with an unidentified transmission route [77, 78]. Among Enterobacteriaceae, the *Arsenophonus* genus differs, in that all strains to date are believed to be arthropod host-restricted and previously environmental acquisition has not been shown to occur routinely [27].

Given that vertical transmission is not operating in the honey bee – *Arsenophonus* association, the questions arises: what are the main drivers of horizontal transmission? Do honey bees acquire infection predominantly from infected conspecifics, or are infections acquired from the wider environment (e.g. additional host species or reservoirs)? While the symbiotic phenotype of the honey bee – *Arsenophonus* remains unknown, we identified horizontal (infectious) transmission when general contact between conspecifics was allowed, and when contact was via trophallaxis exclusively. For a heritable symbiont, maintenance or reversal to the ancestral (horizontal) transmission route has a clear adaptive benefit in a eusocial host, typified by high host density and low genetic diversity [79], and social acquisition can prevent workers being the evolutionary dead ends they are often considered for heritable symbionts [80]. Localisation in the gut and detection in faeces is consistent with a faecal – oral route, which could allow effective social transmission within a colony and to other hosts via shared floral resources [81–83]. Indeed, *Arsenophonus* has previously been detected on flowers [84], but until now direct evidence for the potential capacity to transmit via this route was lacking.

The marked seasonal incidence of *Arsenophonus*, peaking in early autumn, has a number of potential drivers. Shedding and accumulation of the bacterium in the foraging environment (e.g. on flowers) may generate increased infection risk as the foraging season proceeds, as observed for some parasite species [85]. However this would be contingent on the survival of *Arsenophonus* in the environment, a trait conventionally considered absent from this class of insect symbionts, despite relatively large symbiont genomes and cell free cultivability found within the clade [86, 87]. Alternatively, but not mutually exclusively, *Arsenophonus* dynamics may reflect the activity period of an environmental reservoir or additional host species that is responsible for transmission to honey bees. Comparable seasonality is observed for *Spiroplasma* in honey bees, with peak prevalence of *S*. *melliferum* aligning with peak flowering periods [88, 89]. This latter hypothesis may explain our observation that apiary is a significant predictor of a colonies *Arsenophonus* status, while over broad spatial scales infection prevalence does not vary notably. Here, localised spatial pattern reflects shared exposure to reservoirs of infection, with colonies originating from the same apiary often overlapping in foraging area [90]. Relevant here are reports of *Arsenophonus* associated with pollen and nectaring sites [42, 84], providing further evidence for environmental transmission [84]. Other processes, such as drifting of infected bees within apiaries, could also drive inter-colonial transmission of *Arsenophonus* and feedback to drive the localised infection prevalence we observed.

The position of the honey bee *Arsenophonus* on the parasitism-mutualism continuum remains speculative. Our observations of HT, systemic infections and a higher prevalence among dead hosts strengthens interpretations of previous work correlating *Arsenophonus* with poor health outcomes in bees [44, 45, 91]. HT is considered to increase the scope for the evolution of virulence [17, 18], although this trait alone is insufficient to draw conclusions regarding symbiont phenotype, as many avirulent or beneficial microbes are transmitted horizontally [15, 92]. The association with dead hosts may implicate *Arsenophonus* directly or represent opportunistic proliferation in a compromised host. Alternatively, saprophytic growth on cadavers may be occurring, a capacity demonstrated by *A*. *nasoniae* in fly puparia [93]. Further work is needed to determine if health outcomes are causal in honey bees and to characterise the transmission phenotype of *Arsenophonus* reported from other bee species [91, 94–96].

To conclude, the honey bee - *Arsenophonus* contrasts in its phenotype from the rest of the heritable clade, as vertical transmission is not the predominant transmission mode. Instead, horizontal transmission occurs via social interactions, colonisation can occur in the gut, and environmental reservoirs appear to be necessary for maintenance of the association. Our data establish the honey bee - *Arsenophonus* as a key link in a symbiont clades trajectory from free-living ancestral life to obligate mutualism threatened by mutational decay. This will provide new avenues for research into the emergence of symbiotic diversity.

## Author contributions

The project was conceived initially by GB, CF and GH. GD designed and completed survey, experimental and statistical analysis, with input from GH and GB. PN, OY and SS completed the sequencing and phylogenomic analysis elements. GD wrote the paper with assistance from all authors.

### Acknowledgements

We thank Dr Katherine Roberts, Dr Laurent Gauthier and Dr Kirsty Stainton for assistance on the project and Dr Andri Manser for statistical input. Jack Wilford, and many other bee keepers, were pivotal in sampling efforts. This work was financially supported by a BBSRC iCASE studentship to GD (BB/L016133/1) and Bee Disease Insurance Ltd. Microscopy was conducted at the Centre for Cell Imaging (CCI), using equipment under grant BB/M012441/1.

## Competing interests

We are not aware of any competing interests.

## Notes

### Competing Interest Statement

The authors have declared no competing interest.

