## Supplementary information for "Transitions in symbiosis: evidence for environmental acquisition & social transmission within a clade of heritable symbionts"

### Supplementary Methods

##### Honey bee maintenance (laboratory conditions)

Unless otherwise stated, under laboratory conditions Western honey bees were maintained on filter paper in plastic deli pot cages at 32°C ± 2 °C and incubators were kept dark. Ambrosia® bee fondant and 50% sucrose solution were fed *ad libitum.*

##### DNA extraction by Promega Wizard® purification

A Promega Wizard® genomic DNA purification kit with a protocol modified for insects was used to extract DNA from honey bee samples. Samples were homogenised in nuclei lysis solution (250μl) and incubated for 30 min at 65°C. Protein precipitation solution (80μl) was added and samples were vortexed and held on ice (5 min). Samples were centrifuged (14,725 rpm, 4 min) and supernatant mixed with filtered isopropanol (150μl) via inversion. After further centrifugation (14,725 rpm, 2 min), supernatants were discarded and 70% EtOH (150μl) added to pellets, before gentle vortexing and centrifugation (14,800rpm, 1 min). Supernatants were discarded and pellets airdried (65°C, 30 min) before resuspension in molecular grade $H_20$ (60µl) at 4°C overnight.

#### PCR cycling & sequencing conditions

Polymerase chain reaction assays (PCR) were based on a total volume of 15µl, containing GoTaq® Hot Start Green Master Mix (7.5µl), nuclease free water (5.5µl), forward and reverse 10µM primer (0.5µl) and template DNA (1µl). PCR amplifications were performed on Applied Biosystem® Veriti cycler under the following conditions: 95°C for 2 min, 35 cycles at 94°C for 30 s, 58°C for 30 s (variable, depending on primer Tm) 72°C for 1 min with a final extension of 72°C for 10 min. Primers used for detection of *Arsenophonus* targeted *fbaA* (5' -GCCGCTAAGGTTGGTTCTCC – 3' and 5' - CCTGAACCACCATGGAAAACAAAA – 3') and were adapted from a previous study [1]. Products were visualised on 1.5% w/v agarose gel with 0.5µg/mL ethidium bromide (110 v for 50 min) Every run included a positive, negative and no template control. Where samples were to be sequenced, PCR products were cleaned of unincorporated primers and nucleotides using 0.2µl of SAP (shrimp alkaline phosphatase), 0.05µl of Exonuclease I, 0.7µl of 10X RX Buffer and 1.05µl of molecular grade $H_20$ and incubated at 37°C for 45 min, followed by 80°C for 15 min. Purified products were Sanger sequenced through both strands in house on an ABI® Prism 3010x or outsourced to GATC Biotech.

#### Validation of *Arsenophonus* PCR assay sensitivity

Sensitivity of the detection method for *Arsenophonus* was assessed by serial dilution of *Arsenophonus* template with uninfected honey bee template DNA. In all tested cases (N = 6) *Arsenophonus* DNA remained detectable at $10^{-1}$ and $10^{-2}$ dilutions, in 4/6 cases *Arsenophonus* was detectable at $10^{-3}$ and $10^{-4}$ dilutions of honey bee template, providing confidence in the sensitivity of our detection method.

#### Selection criteria for colonies used in infection maintenance & social transmission experiments

As the *Arsenophonus* status of individual bees included in the infection maintenance and social transmission experiment could not be pre-determined (demands destructive sampling), we selected individuals from colonies with a threshold *Arsenophonus* prevalence of 85%. For each colony this was determined by screens of 15 worker bees individually. Samples were washed in sterile $H_20$ and exposed to ultra violet (UV) light for 10 minutes to cross link contaminating external DNA. DNA was extracted using a Promega Wizard® genomic DNA purification kit with a protocol modified for insects and screened for *Arsenophonus* using PCR assays (as described above). Workers used in negative control assays were from colonies where no adult worker had returned a positive result for *Arsenophonus* detection. For the social transmission study, a sample of NEWs (although generally considered sterile [2] and rarely associated with *Arsenophonus*, see main text Figure 5) were similarly screened to ensure *Arsenophonus* was not present prior to mixing with infected conspecifics. For the transmission experiment, 85% and 0% were thus taken as a proxy starting prevalence (dotted lines on main text Figure 7) for workers and NEWs respectively.

#### Negative control images for localisation of *Arsenophonus* within the gut

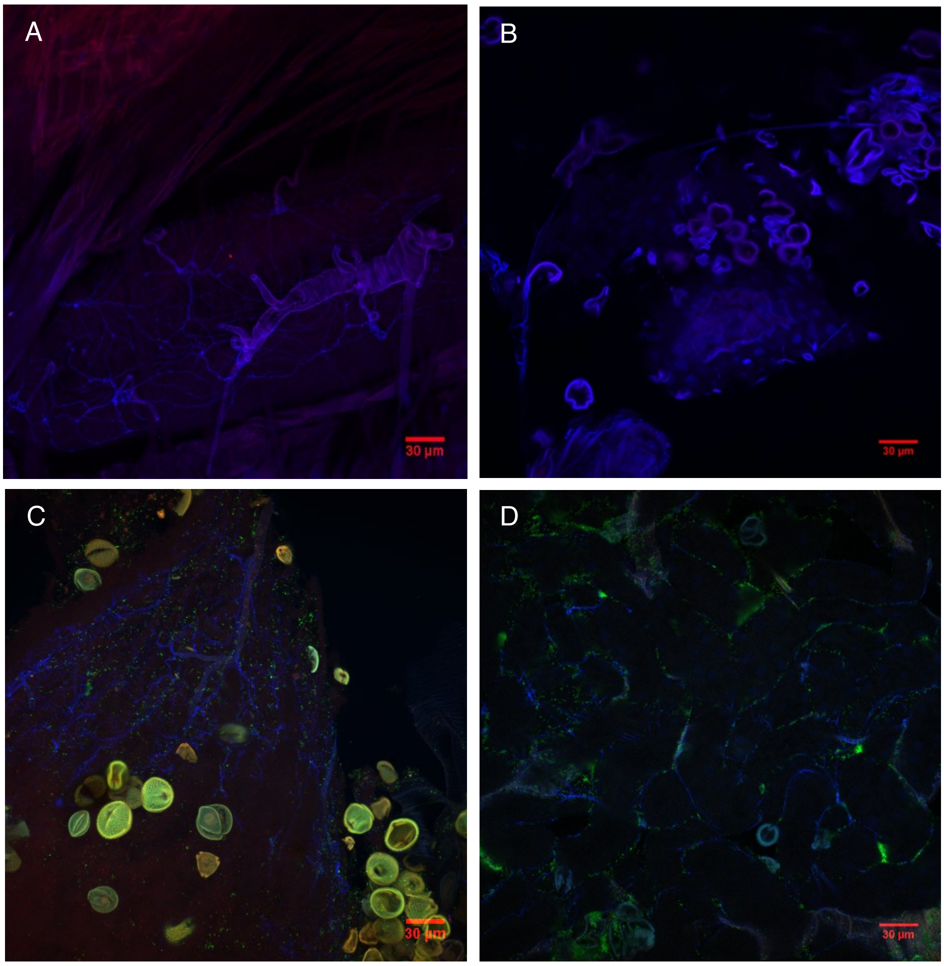


**SI Fig. 1 Confocal microscopy images of whole mounted honey bee guts from colonies with no evidence of *Arsenophonus* association.** Images were obtained using whole gut mounts from *Arsenophonus* negative honey bee workers. These visualisations were used to assess if signals from tissue autofluorescence or inadequate washing would generate non-specific signals from the *Arsenophonus* specific probe (red fluorescence). The absence of significant red signal suggests this was not of concern. Honey bee whole guts (A, B, C, D) were counterstained with DAPI (blue fluorescence) and symbiont DNA targeted by an *Arsenophonus* specific probe (TCATGACCACAACCTCCA) [3] 5' labelled with a Alexa Fluor® 647 fluorochrome (red fluorescence, not visible). A universal bacterial FISH probe is shown highlighting other bacterial members of the gut in green fluorescence (C & D only) (TGCTGCCTCCCGTAGGA) [4], 5' tagged with a Alexa Fluor® 555 fluorochrome. Images were obtained using a ZEISS LSM 880 confocal microscope with a x 40 objective and processed using Image J [5].

#### Selection tables & parameter estimates from statistical models

**SI Tab. 1** Model selection & parameter estimates for spatial & seasonal dynamics of *Arsenophonus* in honey bee colonies

| **Model structure $^{GLMM}$** | ***df*** | **AIC** | **Loglik** | **Dev.** |
|---|---|---|---|---|
| *Arsenophonus* status ~ Day of season$^3$ + (Year) + (County/Apiary/Colony.ID) | 8 | 256.7 | -120.4 | 240.7 |
| *Arsenophonus* status ~ Day of season$^3$ + (Year) + (Apiary/Colony.ID) | 7 | 254.7 | -120.4 | 240.7 |
| *Arsenophonus* status ~ Day of season$^3$ + (Year) + (Apiary) $^{SM}$ | 6 | 252.7 | -120.4 | 240.7 |
| *Arsenophonus* status ~ Day of season$^3$ + (Apiary) | 5 | 254.6 | -122.3 | 244.6 |
| *Arsenophonus* status ~ Day of season$^3$ + (Year) | 5 | 264.5 | -127.3 | 254.5 |

| **Fixed effect** | **Est.** | **SE** | ***Z*** | ***P*** |
|---|---|---|---|---|
| Intercept | -0.7423 | 0.4502 | -1.649 | 0.09920 |
| Day of season$^1$ | 17.37 | 4.514 | 3.849 | 0.000118*** |
| Day of season$^2$ | -12.26 | 4.162 | -2.946 | 0.003223** |
| Day of season$^3$ | -7.736 | 3.495 | -2.213 | 0.02687* |

| **Random effect** | ***N*** | **Var.** | **SD** |
|---|---|---|---|
| Apiary | 45 | 1.155 | 1.075 |
| Year | 5 | 0.4944 | 0.7031 |

Note, day of the season (March 20th = day 0, November 8th = day 232) was modelled as a fixed effect 3rd order polynomial. A total of 229 observations of *Arsenophonus* status were included in the analysis. $^{SM}$ Selected model, $^{df}$ Degrees of freedom, $^{AIC}$ Akaike's information criterion, $^{Dev.}$ Deviance, $^{Est.}$ Coefficient estimate, $^{Var.}$ Variance. (*P* <***0.001, **0.01, *0.05)

**SI Tab. 2** Model selection & parameter estimates for *Arsenophonus* loss in individuals removed from the colony & foraging environment

| Model structure GLMM | *df* | AIC | Loglik | Dev. |
|---|---|---|---|---|
| *Arsenophonus* status ~ Time * Colony + Treatment + (1 \| Pot) | 6 | 100.7 | -44.3 | 88.7 |
| *Arsenophonus* status ~ Time * Colony + (1 \| Pot) | 5 | 98.94 | -44.5 | 88.9 |
| *Arsenophonus* status ~ Time + Colony + (1 \| Pot) SM | 4 | 97.52 | -44.8 | 89.5 |
| *Arsenophonus* status ~ Time + (1 \| Pot) | 3 | 115.8 | -54.9 | 109.8 |
| **Fixed effect** | **Est.** | **SE** | ***Z*** | ***P*** |
| Intercept | 6.661 | 1.384 | 4.813 | < 0.001*** |
| Time | -0.3736 | 0.0739 | -5.055 | < 0.001*** |
| Colony (Colony B) | -3.615 | 1.058 | -3.414 | < 0.001*** |
| **Random effect** | ***N*** | **Var.** | **SD** | |
| Pot | 19 | 0.9725 | 0.9862 | |

SM Selected model, *df* Degrees of freedom, AIC Akaike's information criterion, Dev. Deviance, Est. Coefficient estimate, Var. Variance. (*P* <***0.001, **0.01, *0.05)

**SI Tab.3** Horizontal transmission of *Arsenophonus* in honey bees under two social conditions

| Model structure GLMM | *df* | AIC | Loglik | Dev. |
|---|---|---|---|---|
| *Arsenophonus* status ~ Bee status * Transmission + Colony + Fate + (1\| Pot) | 9 | 307.2 | -144.6 | 289.2 |
| *Arsenophonus* status ~ Bee status + Transmission + Colony + Fate + (1\| Pot) | 8 | 306.2 | -145.1 | 290.2 |
| *Arsenophonus* status ~ Bee status + Transmission + Fate + (1\| Pot) SM | 5 | 303.3 | -146.7 | 293.4 |
| *Arsenophonus* status ~ Bee status + Transmission + (1\| Pot) | 4 | 309.0 | -150.5 | 300.9 |
| *Arsenophonus* status ~ Bee status + Fate + (1\| Pot) | 4 | 306.6 | -149.3 | 298.6 |
| **Fixed effect** | **Est.** | **SE** | ***Z*** | ***P*** |
| Intercept | -1.612 | 0.5545 | -2.907 | 0.00365 ** |
| Status (NEW) | -0.5058 | 0.3298 | -1.534 | 0.1251 |
| Transmission (General contact) | 1.842 | 0.7714 | 2.387 | 0.01697 * |
| Fate (Dead) | 1.930 | 0.7385 | 2.613 | 0.00898 ** |
| **Random effect** | ***N*** | **Var.** | **SD** | |
| Pot | 20 | 2.371 | 1.54 | |

Bee status = Worker *or* NEW, Transmission = Gen contact *or* Trophallaxis, Fate = Dead *or* Alive. Total observations (N = 300). SM Selected model, *df* Degrees of freedom, AIC Akaike's information criterion, Dev. Deviance, Est. Coefficient estimate, Var. Variance. (*P* <***0.001, **0.01, *0.05)
